# Evaluating FoldX 5.1 for MAVISp Stability Data Collection

**DOI:** 10.64898/2026.03.31.715598

**Authors:** Angeliki Vliora, Matteo Tiberti, Elena Papaleo

## Abstract

MAVISp (Multi-layered Assessment of VarIants by Structure for proteins) is a structure-based framework for facilitating mechanistic interpretation of missense variants, with protein stability as one of its core analytical layers. When software tools are updated, a key consideration for database curation is whether the new version can be adopted without compromising compatibility with existing entries. This study evaluated the effect of replacing FoldX5 with FoldX5.1 on the results of the MAVISp stability workflow. We compared predicted changes in folding free energy for 539,809 shared variants across 119 proteins. We found high overall agreement with a mean Pearson correlation of 0.933 and a mean Cohen coefficient of 0.814. Most proteins showed strong concordance. The number of disagreements was higher at sites with low AlphaFold2 confidence. These outliers did not display systematic inter-version bias, as mean shifts in folding free energies between versions were minimal. Collectively, these findings support adopting FoldX5.1 for future MAVISp data collection. We will include a transition period, during which existing entries retain FoldX5 annotations until their scheduled annual update, while new or updated entries are processed with FoldX5.1. To facilitate this transition, the FoldX software version has been added as a new metadata annotation in the MAVISp database.

## Introduction

Computational frameworks need to balance methodological improvement with database continuity [1]. We recently released MAVISp [2] as a structure-based framework for evaluating variant effects on protein structure using multiple mechanistic indicators. In MAVISp, predicted effects on protein stability are one of the key mechanistic layers used to contextualize missense variants. We aim to maintain and update the database regularly, both considering the main dataset and the underlying methodology. As MAVISp is updated iteratively with each release, replacing a well-estabilished component of the stability prediction workflow with a more recent version raises practical questions: should the framework continue to rely on a legacy tool version for consistency and delay adoption until all entries are recomputed, or allow temporary coexistence of legacy and updated calculations?

In this work, we directly addressed this issue for FoldX [3,4]. FoldX is a widely used empirical force-field method for predicting the effect of amino acid substitutions on protein structural stability and binding, and it is commonly applied in large-scale analyses [2,5–8]. Its computational efficiency stems from local optimization around the mutated residue, without backbone rearrangements. While FoldX is currently implemented as a single-core executable, Python bindings or wrappers, such as pyFoldX [9] and Mutatex [10], allow parallelizing several FoldX runs, taking advantage of modern multi-core systems. The newer FoldX 5.1 [11] version maintains the same overall framework as the previous versions, but includes several force-field updates, such as pi-stacking terms, broader pH dependence for charged residues, improved aromatic interaction terms, revised helix-related contributions, adjustments to hydrogen-bonding, side-chain entropy, and solvation parameters, all introduced to improve performance against experimental stability data.

In MAVISp, the STABILITY module employs multiple predictors and a consensus classification scheme to infer whether each mutation affects protein stability [2], including FoldX. In this contribution, we have followed the MAVISp protocol for high-throughput data collection (*simple mode*) for FoldX, collected stability predictions using either FoldX 5 or 5.1, and compared the two datasets for a set of 119 representative proteins, chosen to capture a heterogeneous sample of several protein folds. We restricted this comparison to the MAVISp *simple mode* (i.e., using a single structure per protein) to better represent a large-scale, high-throughput collection scenario, which is used for most entries currently available in MAVISp. We show that results obtained using FoldX 5.1 closely resemble those obtained with FoldX 5 across most proteins currently assessed, that discrepancies are concentrated in a small number of structurally uncertain cases, or can be related to improvements in the FoldX 5.1 energy function. Overall, the available evidence supports a phased replacement of FoldX 5 in the MAVISp data curation.

## Results and Discussion

### Construction of the comparison dataset

We first constructed two datasets to compare predictions from FoldX5 and FoldX5.1. To do so, we selected a set of representative proteins spanning a heterogeneous range of folds and protein structures to limit structure-specific bias.

Our dataset included 119 proteins present in the MAVISp database as of 20/05/2025, selected by removing redundancy in protein folds across the entire MAVISp database (see Methods). We then collected data on each of these proteins using FoldX 5.1, for a total of 539,809 analyzed variants, and used FoldX 5 data already available in the MAVISp dataset.

It should be noted that the starting structures used in the FoldX 5 and FoldX 5.1 datasets differed in a few ways, as described in the next section. We therefore sought to quantify the differences in starting structures between the two datasets.

### The structure models used with FoldX5 and FoldX5.1 are mostly similar, with a few exceptions

Each of the FoldX 5 and FoldX 5.1 datasets was independently collected by means of an automated pipeline as per the standard MAVISp protocols (see Methods for details), which downloads protein structure models from the AlphaFold Protein Structure Database (AFPSD) [12] and trims them according to user-specified residue ranges, in order to isolate structured domains and leave out of the analysis protein regions that are either fully unstructured or with a low prediction confidence, as evidenced by their pLDDT score. As the FoldX 5 and FoldX 5.1 datasets were collected at different times, neither the selected trimming strategy nor the structural models used were always the same between the two. Furthermore, the FoldX 5 dataset included proteins previously annotated in the MAVISp database for which a model produced using AlphaFold 3 [13] was used; these were CNOT3, DEFB108B, MLF1 and ZNF738.

For most of the proteins we considered, the protein-structure regions selected as input for prediction (trimming) were identical across the FoldX5 and FoldX5.1 datasets. Trimming varied among 17 proteins: 11 had differences of just 1 residue at the N-terminus, and the remaining 6 showed larger differences (**Supplementary Information Table S1**). In the following analyses, we have considered only mutations at positions present in both the FoldX 5 and FoldX 5.1 datasets.

Furthermore, because the two datasets were produced at different times and due to updates at the AFPSD, protein structures in the two datasets didn’t necessarily have identical conformations. Therefore, we pairwise compared the initial structures used for the FoldX 5 and FoldX 5.1 calculations using C_α_ RMSD (see Methods) after superimposition. The calculated RMSD values were below 5 Å (median 0.4 Å) for most proteins (**Supplementary Material Text S1**). A few of them featured higher RMSD values, in the ∼6-21.7 Å range: these were ZNF738, ADCK2, SMARCE1, RSPO2, CBX8, ASXL2, FANCE, NUPR1, and CBFA2T3. Visual inspection (Supplementary Material Text S1) showed that the differences were system-dependent. For ZNF738 (RMSD 21.7 Å), the overall conformation differed significantly between the two cases; for RSPO2 (RMSD 11 Å), the positioning of a terminal α-helix differed between the two models; for NUPR1 (RMSD 7.7 Å), the conformations of the two models also differed markedly. For the remaining cases, differences could be attributed to conformational changes in flexible loops, which were modelled with low confidence.

### Global agreement between FoldX versions across the shared MAVISp set

We observed strong agreement across the benchmark set, with an overall Pearson correlation coefficient of 0.956 and a mean absolute error of 0.52 kcal/mol (**Figure 1A-B)**. We have also compared datasets on a per-protein basis, yielding a mean Pearson correlation of 0.933 ± 0.077 and a mean Cohen’s κ of 0.814 ± 0.092 (**see OSF for data on individual proteins; Figure 1C-D**).

**Figure 1.**
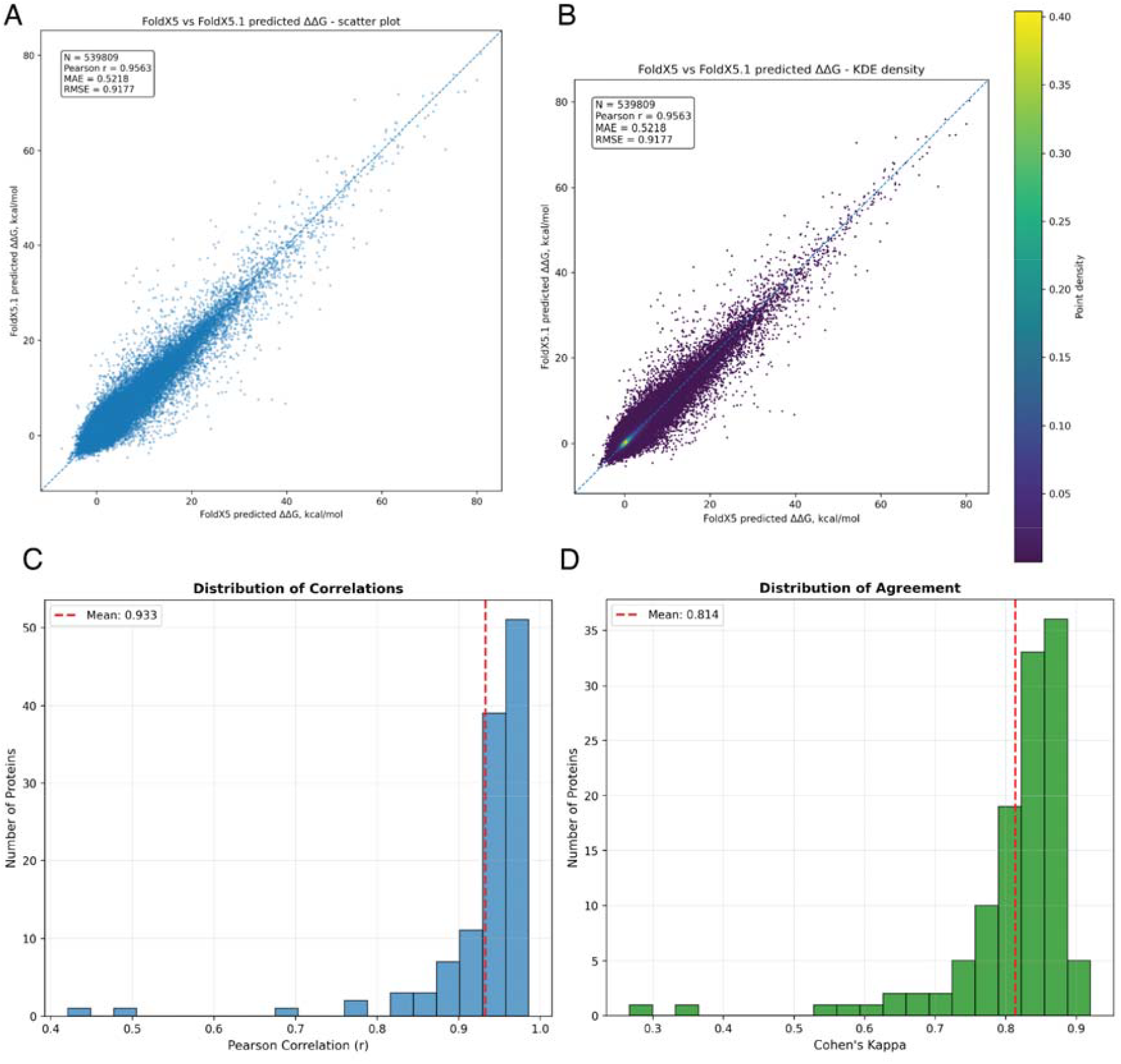
Agreement between the FoldX 5 and FoldX 5.1 dataset. (A) scatter plot of predicted ΔΔG values for FoldX5 and FoldX5.1 considering the whole dataset. (B) The same data from panel A were used to compute a kernel density estimate, which was then sampled at the data points. This was done to visualize the data density, given the large number of data points. ( C) Distribution of Pearson correlation coefficient values, each calculated from the FoldX and FoldX5.1 ΔΔG datasets for a single protein. (D) Distribution of Cohen’s κ coefficient over our dataset, each calculated from the FoldX and FoldX5.1 ΔΔG datasets for a single protein.

A few proteins in the dataset were characterized by a lower agreement than the rest. These proteins were then flagged for follow-up if they met at least one of three criteria: Pearson correlation r < 0.7, Cohen’s κ < 0.5, or classification accuracy < 75% (**see Methods**). Based on these criteria, three proteins required further attention: TMEM127, NUPR1, and TSC1 (**Figure 2**). NUPR1 showed r = 0.491 and κ = 0.267 across 1121 shared variants. TSC1 showed r = 0.680 and κ = 0.532 across 2161 shared variants. Finally, TMEM127 featured r = 0.421 and κ = 0.349 across 475 variants.

**Figure 2.**
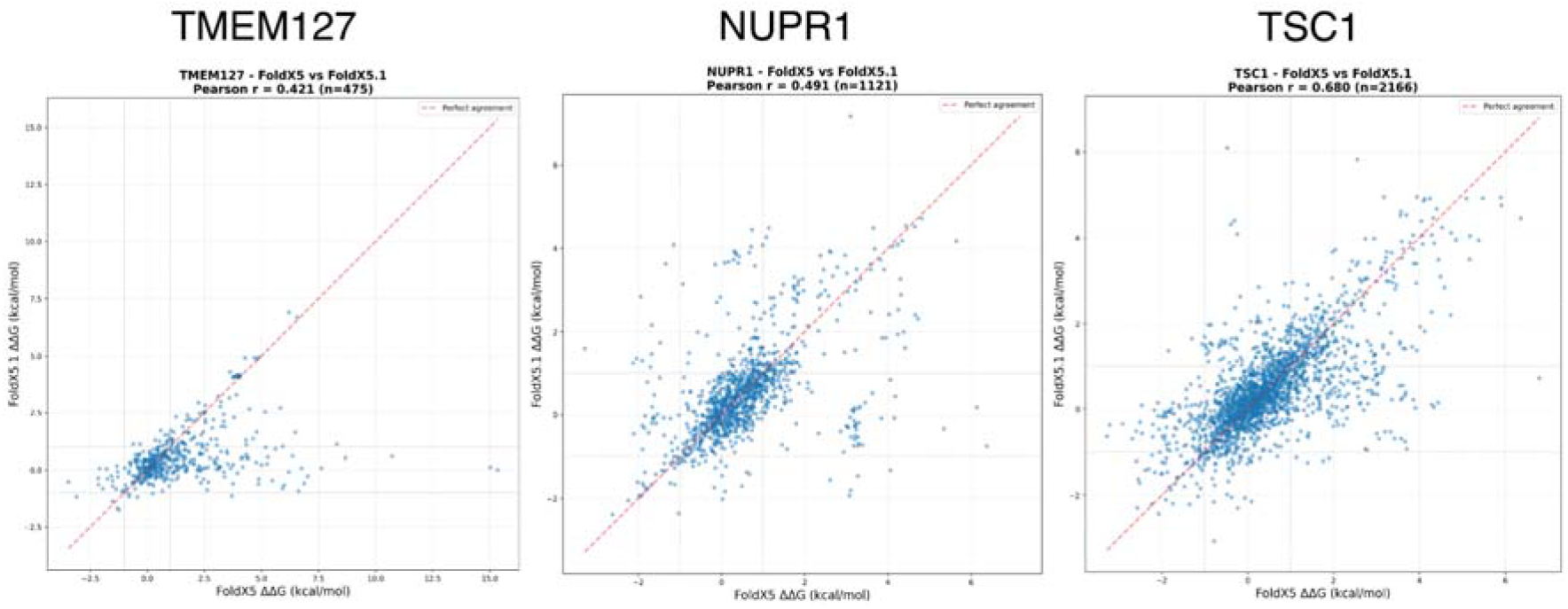
Predicted ΔΔG values for specific proteins. Scatter plots of predicted ΔΔG values for FoldX5 and FoldX5.1 for proteins TMEM127, NUPR1 and TSC1

### Sites with disagreements are enriched in structurally uncertain regions for some proteins, or connected to differences in structural models

To assess whether outlier behavior reflected intrinsic differences in methodology or was due to input structure quality, AlphaFold2 pLDDT values at mutated positions were compared across all analyzed sites, excluding those proteins for which pLDDT scores were not available in the MAVISp dataset (CNOT3, DEFB108B, FUS, GPC3, MLF1, ZNF738). We considered the MAVISp classification for FoldX 5 and FoldX 5.1 data and evaluated whether it was the same. We stratified variants by their agreement or disagreement in regards to MAVISp classification, using either FoldX5 or FoldX5.1, and compared the pLDDT distributions of the two populations. Our global analysis showed that the agreement and disagreement classes have similar distributions, suggesting that cases where the two methods differ are not particularly enriched in low-quality or unstructured regions (**Figure 3A**). Furthermore, just ∼11% of variants were classified differently when using predictions from the two versions, consistently with a broad agreement between the two, as previously discussed.

**Figure 3.**
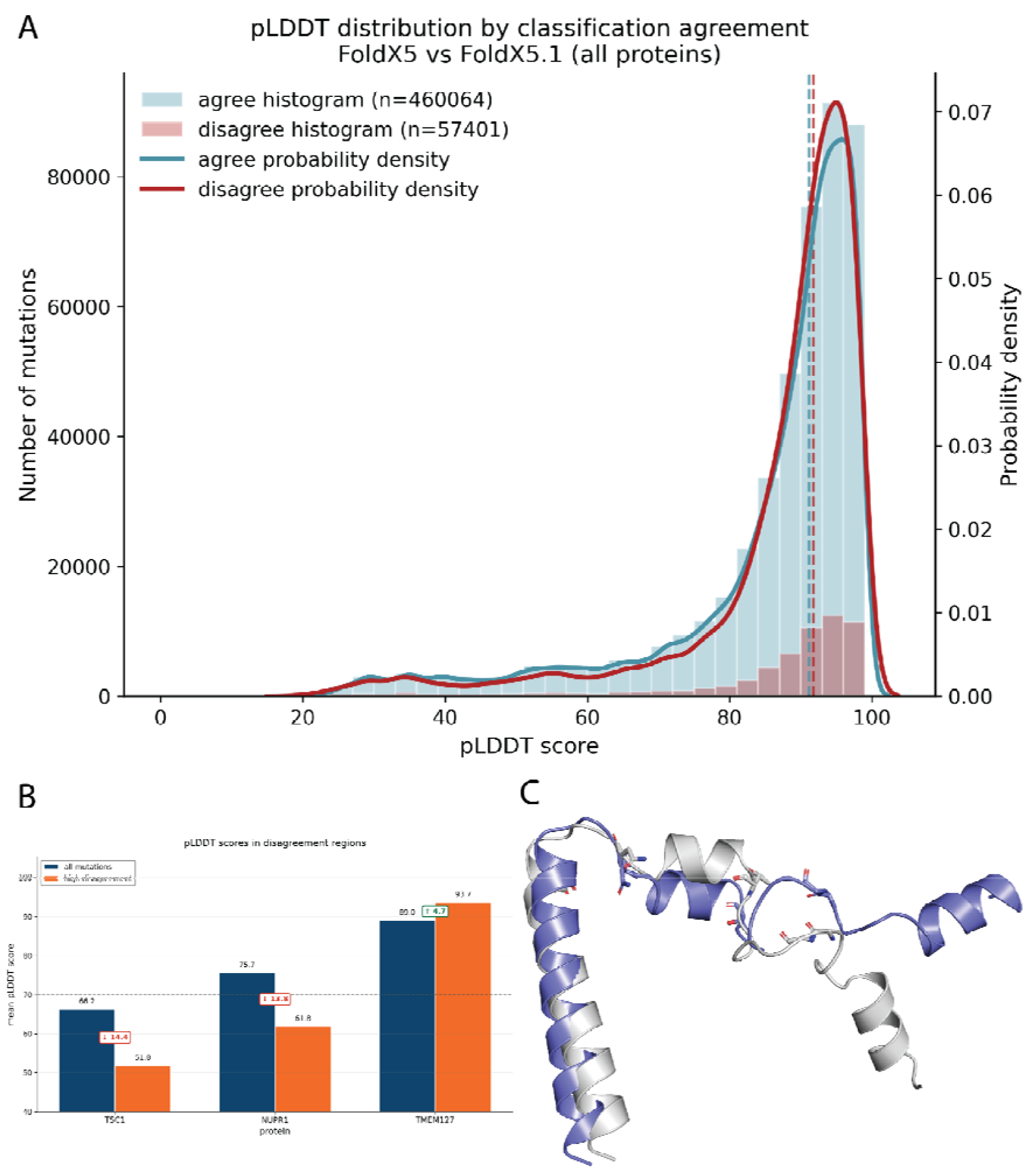
Relation between structure quality and classification agreement. **(A)** Distribution of pLDDT scores for the agreement (same classification) and disagreement (different classification) classes in our dataset. It should be noted that the number of data points in this plot is different than for Figure 1, because of missing data in our dataset, as discussed in the main text. Dashed lines show the median value for each distribution **(B)** Average pLDDT values for either all mutation sites or mutation sites on which mutations were found to be classified differently using FoldX 5 or FoldX 5.1, for a selection of proteins. **(C)** Protein structures of NUPR1, showing the models used for FoldX 5 (light grey) and FoldX 5.1 (blue). Residue in sticks are G40, T46, N55, and G61, left to right.

Nonetheless, in both NUPR1 and TSC1, the disagreement sites had, on average, lower predicted quality than the overall set of mutation sites. In NUPR1, the mean pLDDT decreased from 75.7 across all analyzed positions to 61.8 at sites of disagreement. In TSC1, the mean pLDDT decreased from 66.2 to 51.8. In both cases, the mean pLDDT of disagreement-associated positions fell below the commonly used pLDDT = 70 confidence threshold for structural quality. A similar pattern was not observed for TMEM127, where the disagreement sites were in a region with high average pLDDT (**Figure 3B**). As previously detailed, we also used RMSD to compare structures used for FoldX5 and FoldX 5.1. For TMEM127, we found very small differences (C_α_ RMSD of 0.4 Å) and similar pLDDT patterns when comparing the structures from the two methods. TMEM127 is a transmembrane protein [14], and the lower agreement observed for this target may reflect, at least in part, the known difficulty of applying general structure-based ΔΔG predictors to membrane proteins, especially if variants in transmembrane segments were retained in the analysis [15]. For TSC1_601-714_, we observed a higher but still reasonable C_α_ RMSD of 3 Å, with most differences in the C-terminal region or loops (**see OSF repository**). On the contrary, for NUPR1_24-82_, as previously described, the two models adopt different conformations despite showing similar pLDDT profiles (**Figure 3C; OSF repository**), with a main-chain RMSD of 6.8 Å. This region is close to the lower limit of the protein inclusion criteria in the MAVISp *simple mode*. It corresponds to a short three-helix fragment with very limited tertiary contacts, so stability predictions should be interpreted with caution. Some disagreement sites, including G40 and T46, occupy different local environments in the two starting models, making it difficult to separate structural-model effects from effects due to differences in the FoldX version. We therefore interpret the NUPR1 discrepancies as arising from a combination of model uncertainty and, for some variants, differences introduced by the revised FoldX5.1 energy function.

More broadly, these results are consistent with the known dependence of structure-based ΔΔG predictors on the quality of the input model [5,16,17].

### Top sites with predictions in disagreement are enriched for amino acid substitutions to aromatic residues or proline

We then examined the variants with the largest inter-version disagreements and found that many involved substitutions to aromatic residues or, in some cases, to proline (**Table 1**). For several of these variants, FoldX5 predicted marked destabilizing effects, whereas FoldX5.1 predicted neutral or less severe effects **(Table 1**). For aromatic substitutions, this pattern is consistent with the revised version of the FoldX 5.1 force field, which introduced updated π- stacking and aromatic interaction terms. For proline substitutions, however, the interpretation is less direct. Still, it could be related to the corrections introduced for the minimum buried volume in FoldX 5.1 or indirect effects of the updated hydrogen-bond and helix-related terms [11], or to sensitivity to the starting structure. Other outlier cases involved mutations at wild-type glycine positions. We do not interpret this as evidence for a glycine-specific change in FoldX 5.1. FoldX 5.1 revised N-cap, α-helix dipole, and hydrogen-bond terms could have an indirect influence on this result. So, if a glycine site sits in or near one of those structural contexts, some differences could also arise indirectly from those terms rather than from glycine itself.

**Table 1.** Top 10 variant outliers in the comparison between FoldX5 and FoldX5.1 for TMEM127, NUPR1, and TSC1. The changes in classification within the MAVISp protocol, using the consensus approach based on FoldX and RaSP, are reported, along with the predicted changes in folding free energy upon amino acid substitutions.

| VARIANT | FoldX5 (kcal/mol) | Foldx5.1 (kcal/mol) | Classification Change using FoldX/RaSP consensus (MAVISp) |
| --- | --- | --- | --- |
| TMEM127 |  |  |  |
| V169W | 15.35 | 0.01 | Uncertain-> Neutral |
| A174H | 15.02 | 0.12 | Destabilizing -> Uncertain |
| V169Y | 10.72 | 0.58 | Uncertain -> Neutral |
| V171W | 8.68 | 0.51 | Destabilizing -> Uncertain |
| V169F | 7.63 | 0.08 | Uncertain-> Neutral |
| A150W | 6.62 | -0.93 | Destabilizing -> Uncertain |
| Q158W | 6.87 | -0.58 | Uncertain -> Neutral |
| A174P | 7.03 | -0.27 | Destabilizing -> Uncertain |
| V169R | 8.31 | 1.10 | Uncertain -> Neutral |
| A174W | 5.92 | -0.81 | Destabilizing -> Uncertain |
| NUPR1 |  |  |  |
| G40P | 6.38 | -0.75 | Destabilizing -> Uncertain |
| N55P | 6.14 | 0.17 | Destabilizing -> Uncertain |
| G39P | 5.35 | -0.34 | Destabilizing -> Uncertain |
| G39I | 4.06 | -1.32 | Uncertain -> Neutral |
| G61H | -1.15 | 4.08 | Neutral -> Uncertain |
| G39F | 3.08 | -1.93 | Uncertain -> Neutral |
| G61W | -1.34 | 3.63 | Neutral -> Uncertain |
| G39Y | 3.11 | -1.80 | Uncertain -> Neutral |
| T46D | -3.28 | 1.60 | Uncertain -> Neutral |
| T46E | -1.94 | 2.84 | Neutral |
| TSC1 |  |  |  |
| P670W | -0.47 | 6.09 | Neutral -> Uncertain |
| G639P | 6.79 | 0.72 | Destabilizing -> Uncertain |
| P670F | -0.31 | 4.39 | Neutral -> Uncertain |
| G680I | -0.39 | 4.31 | Uncertain |
| D605P | 3.71 | -0.93 | Destabilizing -> Uncertain |
| V620W | 3.66 | -0.76 | Uncertain -> Neutral |
| P670Y | -0.24 | 4.08 | Neutral -> Uncertain |
| G639T | 4.34 | 0.05 | Uncertain |
| G639V | 4.44 | 0.05 | Uncertain |
| F679I | 3.57 | 4.27 | Uncertain |

### There is no evidence of systematic inter-version changes

The analysis of the distribution of predicted ΔΔG values for the three main outliers did not reveal a systematic recalibration across methodologies. Mean shifts were minimal to modest, in the range of -0.8 to −0.04 kcal/mol (**Table 2**), and the variance in the prediction remained similar between FoldX 5 and FoldX 5.1. Thus, the observed disagreements appear to be scattered across individual variants rather than driven by a global change in the score scale or a tendency toward destabilization.

**Table 2.**
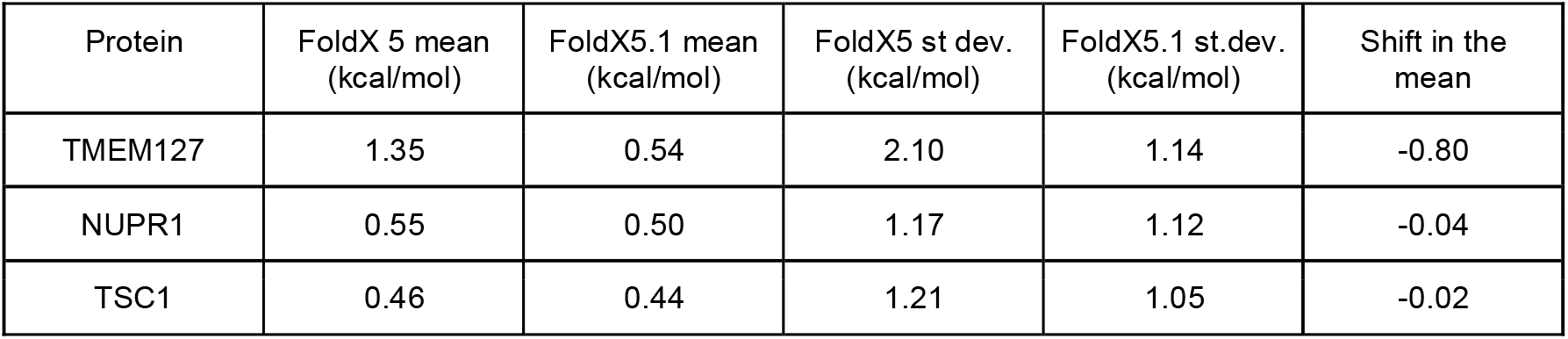
Comparison of the overall ΔΔG distributions predicted by FoldX5 and FoldX5.1 for the three main outlier proteins. For each protein, the table reports the mean and standard deviation of the predicted ΔΔG values across all shared variants, together with the shift in the mean between FoldX 5.1 and FoldX 5. The small changes in the mean indicate that a systematic inter-version shift in predicted stability values does not explain the disagreements observed for these proteins.

### Implications for version migration in MAVISp

Overall, the results support replacing FoldX 5 with FoldX 5.1 in MAVISp stability calculations. For a database updated on an annual schedule, the most practical solution is not to require that all legacy values need to be recomputed before adoption, but to allow the version transition to proceed in phases. Under this model, pre-existing entries can retain their FoldX 5 annotations until their next scheduled update, whereas new or revised entries can be curated with FoldX 5.1. The available evidence suggests that this temporary coexistence is unlikely to compromise interpretation for most proteins, provided that the software version used is explicitly documented [18,19] and structurally uncertain sites are flagged.

## Methods

FoldX version compatibility was assessed in the MAVISp simple-mode stability workflow [2] (https://github.com/ELELAB/mutatex_pipelines/). The comparison was restricted to the 119 proteins for which FoldX data were originally available in the MAVISp database and selected using the *pfam_scrape* workflow [20], which is available in the MAVISp accessory tools repository (https://github.com/ELELAB/mavisp_accessory_tools) and based on Pfam definition of domains [21]. The list of retained proteins is provided as **Supplementary Table S1**. For each protein, only mutations present in both versions were retained, and positions with missing values in either dataset were excluded, yielding 539,809 shared variants.

Our workflow downloads protein structure models from the AlphaFold Protein Structure Database [12] and trims them according to user-specified residue ranges to isolate structured domains and exclude protein regions that are either fully unstructured or low-quality, as evidenced by their pLDDT scores. More details are available in the MAVISp manuscript [2] and website (https://services.healthtech.dtu.dk/services/MAVISp-1.0/).

Quantitative agreement between versions was evaluated by calculating Pearson correlation coefficients for the dataset as a whole and for each protein separately. For the agreement assessment, mutations were classified into three categories using the outer MAVISp classification threshold of ±3.0 kcal/mol. This simplified scheme collapses the MAVISp Uncertain category (−3 < ΔΔG ≤ −2 kcal/mol and 2.0 ≤ ΔΔG < 3.0 kcal/mol into the Neutral class: Stabilizing: ΔΔG < −3.0 kcal/mol, Neutral: −3.0 ≤ ΔΔG ≤ 3.0 kcal/mol, and Destabilizing: ΔΔG > 3.0 kcal/mol. Classification agreements were summarized with raw accuracy and Cohen’s κ (**see OSF repository**).

Proteins were considered poor-agreement cases if they met any of the following criteria: Pearson correlation r < 0.7, Cohen’s κ < 0.5, or classification accuracy < 75%. For these proteins, additional analyses were performed to distinguish genuine disagreement due to methodological differences between the two versions from limitations in structural quality or otherwise.

AlphaFold2 pLDDT values, extracted from the FoldX 5.1 dataset, were compared between all mutated positions and positions at which FoldX 5 and FoldX 5.1 assigned different classes. Comparisons using the pLDDT score are limited to cases where the pLDDT values were available in the MAVISp database, as detailed in Results.

Structure models generated with FoldX 5 or FoldX 5.1 were pairwise compared using Root Mean Square Deviation (RMSD), calculated over all C_α_ atoms of each protein after least-squares superimposition on the same set of atoms. RMSD was calculated using the MDAnalysis Python package [22].

In the underlying MAVISp workflow, FoldX calculations were run through MutateX [10] after structure repair and minimization using the RepairPDB command, followed by FoldX BuildModel-based in silico saturation mutagenesis [23]. The comparison, therefore, reflects version behavior in the exact practical setting relevant to routine MAVISp stability data collection.

## Conclusions

Predictions obtained using FoldX 5.1 showed good agreement with those collected using FoldX 5 and will be adopted for future data collection in MAVISp, without immediately re-evaluating stability data for existing entries in the database. Our analyses displayed a substantial agreement between the two FoldX versions, including 119 proteins and more than half a million shared variants, both in terms of inter-dataset correlation and following MAVISp classification. A small number of proteins displayed a substantial disagreement, which can be explained largely by structural uncertainty at the mutation site or by differences in the definition of the energy functions for aromatic residues, rather than by a general shift in prediction behavior.

At the same time, one should keep in mind that compatibility does not mean perfect interchangeability for every target. Proteins with retained long disordered regions or low-confidence AlphaFold segments warrant extra caution, but these are generally flagged in downstream analysis by the MAVISp framework. Finally, any coexistence of legacy and updated data should remain transparent to users through explicit software-version annotations at entry level. For this purpose, we will introduce a new metadata annotation in the MAVISp database indicating the FoldX version used for the calculations.

A complete rebuild of the existing database before adopting FoldX 5.1 would require substantial time and delay methodological updates, whereas mixing versions would reduce provenance and interpretability. A staged transition is therefore the best compromise: retain legacy FoldX 5 values until the next scheduled yearly update, compute new or refreshed entries with FoldX 5.1, and document software version for the stability module.

## Author contributions

*Conceptualization*: EP *Data Curation*: AV, EP *Formal Analysis*: AV *Investigation:* AV, MT, EP *Funding Acquisition:* EP *Methodology*: AV, MT, EP *Project administration:* EP *Resources*: EP *Software*: AV *Supervision*: MT, EP *Validation*: All the co-authors. *Visualization*: AV, MT, EP *Writing – Original Draft:* EP. *Writing – Review and Editing*: All the coauthors.

## Declaration of generative AI and AI-assisted technologies in the writing process

During the preparation of this work, the author(s) used OpenAI ChatGPT 5.4 to improve the language of the manuscript. After using this service, the authors reviewed and edited the content as needed and took full responsibility for the published article.

## Data availability statement

All data and code, including input files and mutatex outputs, are freely available on OSF (https://osf.io/6hrkt/) and in the GitHub repositories listed in the Method Section.

## Acknowledgements

Our research has been supported by Novo Nordisk Fonden Bioscience and Basic Biomedicine (NNF20OC0065262 and NNF0108314) to the E.P. group. The authors would like to thank Eleni Kiachaki and Estzer Toldi for useful scientific discussion.

## Notes

### Competing Interest Statement

The authors have declared no competing interest.

### Summary of Updates

We have included new analyses based on RMSD and pLDDT comparisions on the entire dataset and updated all the figures and text accordingly.

## References

1 Nourbakhsh M, Degn K, Saksager A, Tiberti M, Papaleo E. Prediction of cancer driver genes and mutations: the potential of integrative computational frameworks. Brief Bioinform. 2024;25(2).

2 Arnaudi M, Utichi M, Degn K, Tiberti M, Beltrame L, Krzesińska K, et al. MAVISp: A modular structure-based framework for protein variant effects. Protein Science. 2026;35(5).

3 Guerois R, Nielsen JE, Serrano L. Predicting changes in the stability of proteins and protein complexes: A study of more than 1000 mutations. J Mol Biol. 2002;320(2):369–387.

4 Delgado J, Radusky LG, Cianferoni D, Serrano L. FoldX 5.0: working with RNA, small molecules and a new graphical interface. Bioinformatics. 2019;35(20):4168–4169.

5 Akdel M, Pires DE V., Pardo EP, Jänes J, Zalevsky AO, Mészáros B, et al. A structural biology community assessment of AlphaFold2 applications. Nat Struct Mol Biol. 2022;29(11):1056–1067.

6 Beltran A, Jiang X, Shen Y, Lehner B. Site-saturation mutagenesis of 500 human protein domains. Nature. 2025;637(8047):885–894.

7 Gerasimavicius L, Liu X, Marsh JA. Identification of pathogenic missense mutations using protein stability predictors. Scientific Reports 2020 10:1. 2020;10(1):1–10.

8 Gerasimavicius L, Livesey BJ, Marsh JA. Loss-of-function, gain-of-function and dominant-negative mutations have profoundly different effects on protein structure. Nat Commun. 2022;13(1):3895.

9 Radusky LG, Serrano L. pyFoldX: enabling biomolecular analysis and engineering along structural ensembles. Bioinformatics. 2022;38(8):2353–2355.

10 Tiberti M, Terkelsen T, Degn K, Beltrame L, Cremers TC, da Piedade I, et al. MutateX: an automated pipeline for in silico saturation mutagenesis of protein structures and structural ensembles. Brief Bioinform. 2022;23(3):bbac074.

11 Delgado J, Reche R, Cianferoni D, Orlando G, van der Kant R, Rousseau F, et al. FoldX force field revisited, an improved version. Bioinformatics. 2025;41(2).

12 Bertoni D, Tsenkov M, Magana P, Nair S, Pidruchna I, Querino Lima Afonso M, et al. AlphaFold Protein Structure Database 2025: a redesigned interface and updated structural coverage. Nucleic Acids Res. 2026;54(D1):D358–D362.

13 Abramson J, Adler J, Dunger J, Evans R, Green T, Pritzel A, et al. Accurate structure prediction of biomolecular interactions with AlphaFold 3. Nature. 2024;630(8016):493–500.

14 Flores SK, Deng Y, Cheng Z, Zhang X, Tao S, Saliba A, et al. Functional characterization of TMEM127 variants reveals novel insights into its membrane topology and trafficking. J Clin Endocrinol Metab. 2020;105(9):e3142–e3156.

15 Kroncke BM, Duran AM, Mendenhall JL, Meiler J, Blume JD, Sanders CR. Documentation of an Imperative To Improve Methods for Predicting Membrane Protein Stability. Biochemistry. 2016;55(36):5002–5009.

16 Pak MA, Ivankov DN. Best templates outperform homology models in predicting the impact of mutations on protein stability. Bioinformatics. 2022;38(18):4312–4320.

17 Valanciute A, Nygaard L, Zschach H, Maglegaard Jepsen M, Lindorff-Larsen K, Stein A. Accurate protein stability predictions from homology models. Comput Struct Biotechnol J. 2023;21:66–73.

18 Sandve GK, Nekrutenko A, Taylor J, Hovig E. Ten Simple Rules for Reproducible Computational Research. PLoS Comput Biol. 2013;9(10):e1003285.

19 Wilkinson MD, Dumontier M, Aalbersberg IjJ, Appleton G, Axton M, Baak A, et al. The FAIR Guiding Principles for scientific data management and stewardship. Sci Data. 2016;3(1):160018.

20 Krzesińska K, Degn K, Llorente A, Giannakopoulou E, Tiberti M, Papaleo E. Deciphering Long-Range Effects of Mutations: An Integrated Approach Using Elastic Network Models and Protein Structure Networks. J Mol Biol. 2025;437(20):169359.

21 Mistry J, Chuguransky S, Williams L, Qureshi M, Salazar GA, Sonnhammer ELL, et al. Pfam: The protein families database in 2021. Nucleic Acids Res. 2021;49(D1):D412– D419.

22 Naughton FB, Alibay I, Barnoud J, Barreto-Ojeda E, Beckstein O, Bouysset C, et al. MDAnalysis 2.0 and beyond: fast and interoperable, community driven simulation analysis. Biophys J. 2022;121(3):272a–273a.

23 Van Durme J, Delgado J, Stricher F, Serrano L, Schymkowitz J, Rousseau F. A graphical interface for the FoldX forcefield. Bioinformatics. 2011;27(12):1711–1712.

