## Supplementary material for "Evaluating FoldX 5.1 for MAVISp Stability Data Collection": Text S1

**Text S1. Evaluation of the differences in protein structures used in the FoldX 5 and FoldX 5.1 datasets.**

In the following document, we include a few useful visualizations to understand the differences between protein structures used in the FoldX 5 and FoldX 5.1 datasets. We show a distribution of the calculated C_α_ RMSD between corresponding proteins in the two datasets, as well as a visual comparison of cases with C_α_ RMSD > 5 Å. In the following figures, models from the FoldX 5.1 dataset are shown in blue, while those from the FoldX 5 dataset are shown in orange. Structures were visualized using PyMOL (The PyMOL Molecular Graphics System, Version 3.0, Schrödinger, LLC.), and superimposed using the *align* PyMOL command. The proteins are ordered by C_α_ RMSD, calculated as described in the main text of the paper.

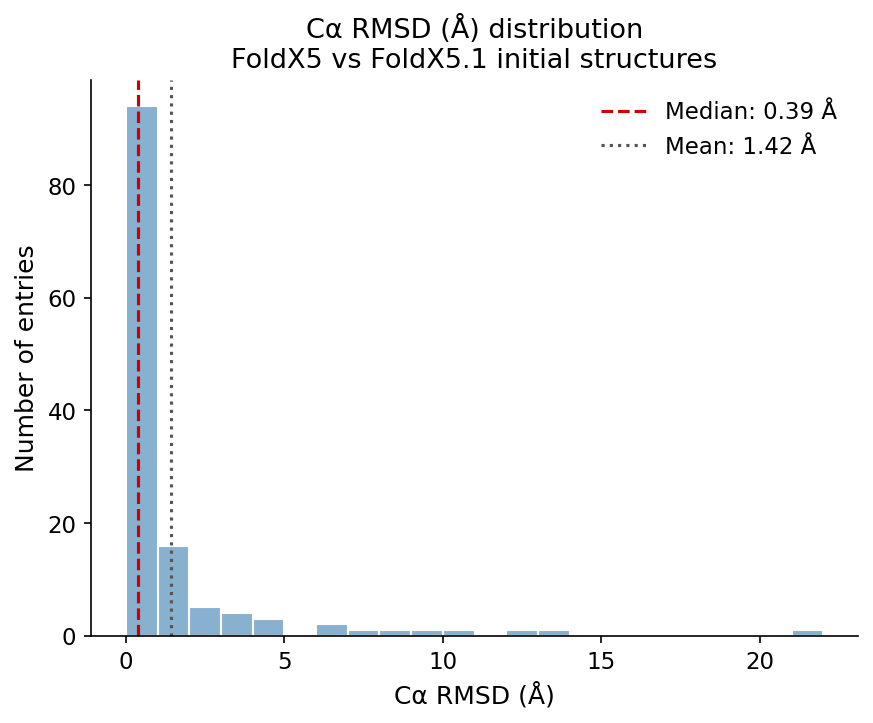

**Figure S1.1:** Distribution of C_α_ RMSD between models used in the FoldX 5 and FoldX 5.1 datasets.

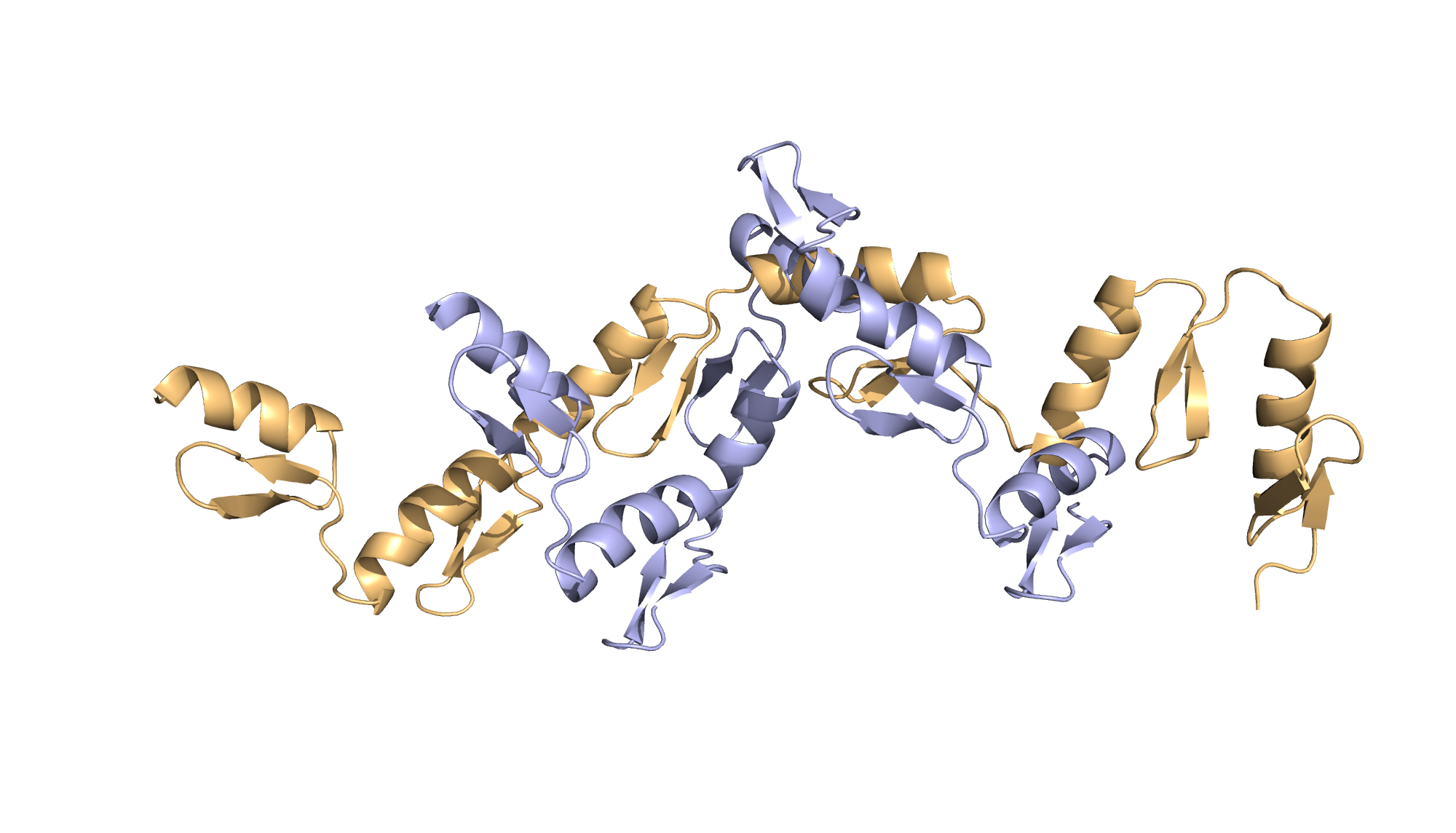

**Figure S1.2:** Visual comparison for ZNF738_201-366_ (C_α_ RMSD 21.7 Å)

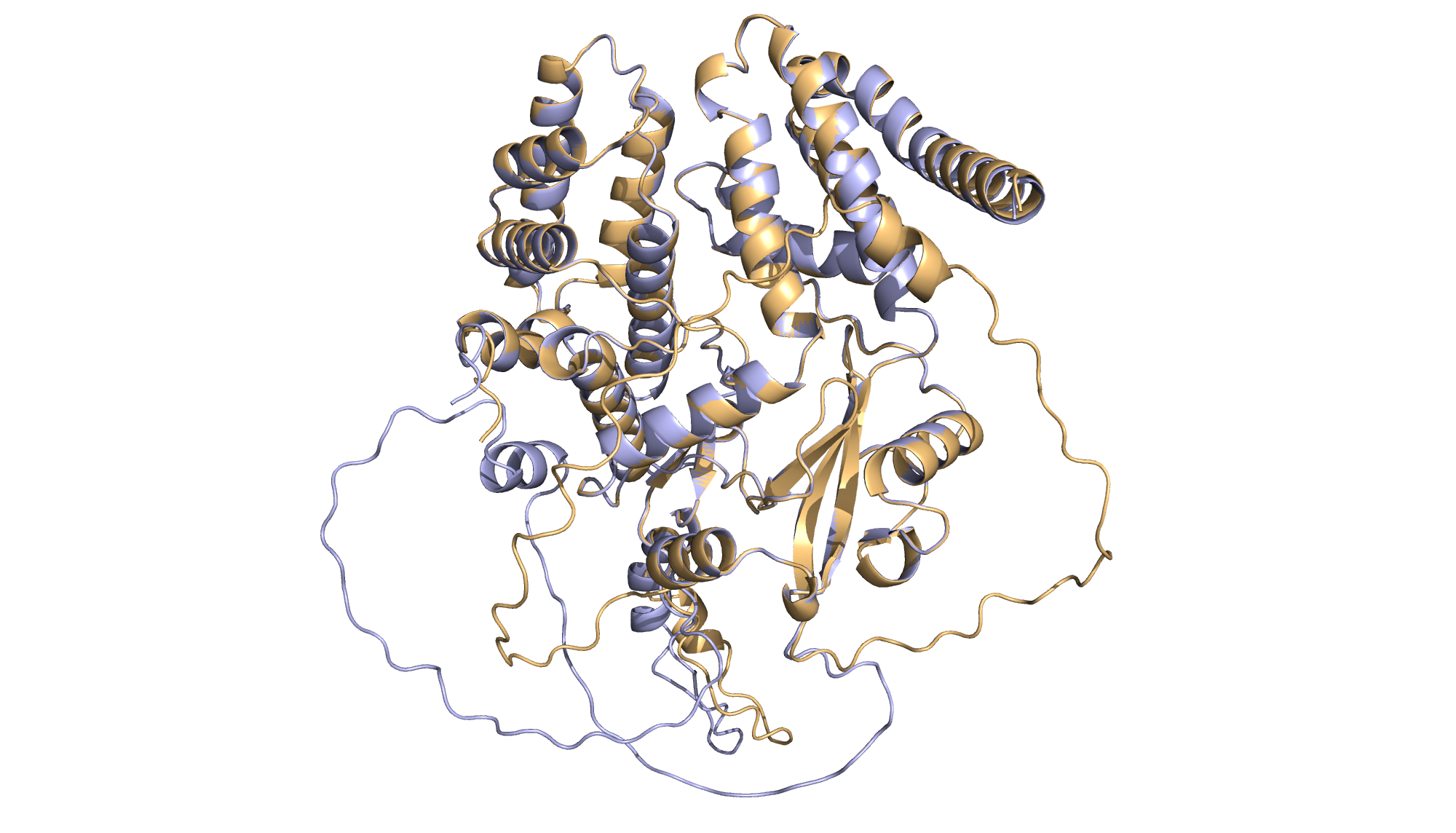

**Figure S1.3:** Visual comparison for ADCK2_88-626_ (C_α_ RMSD 13.4 Å)

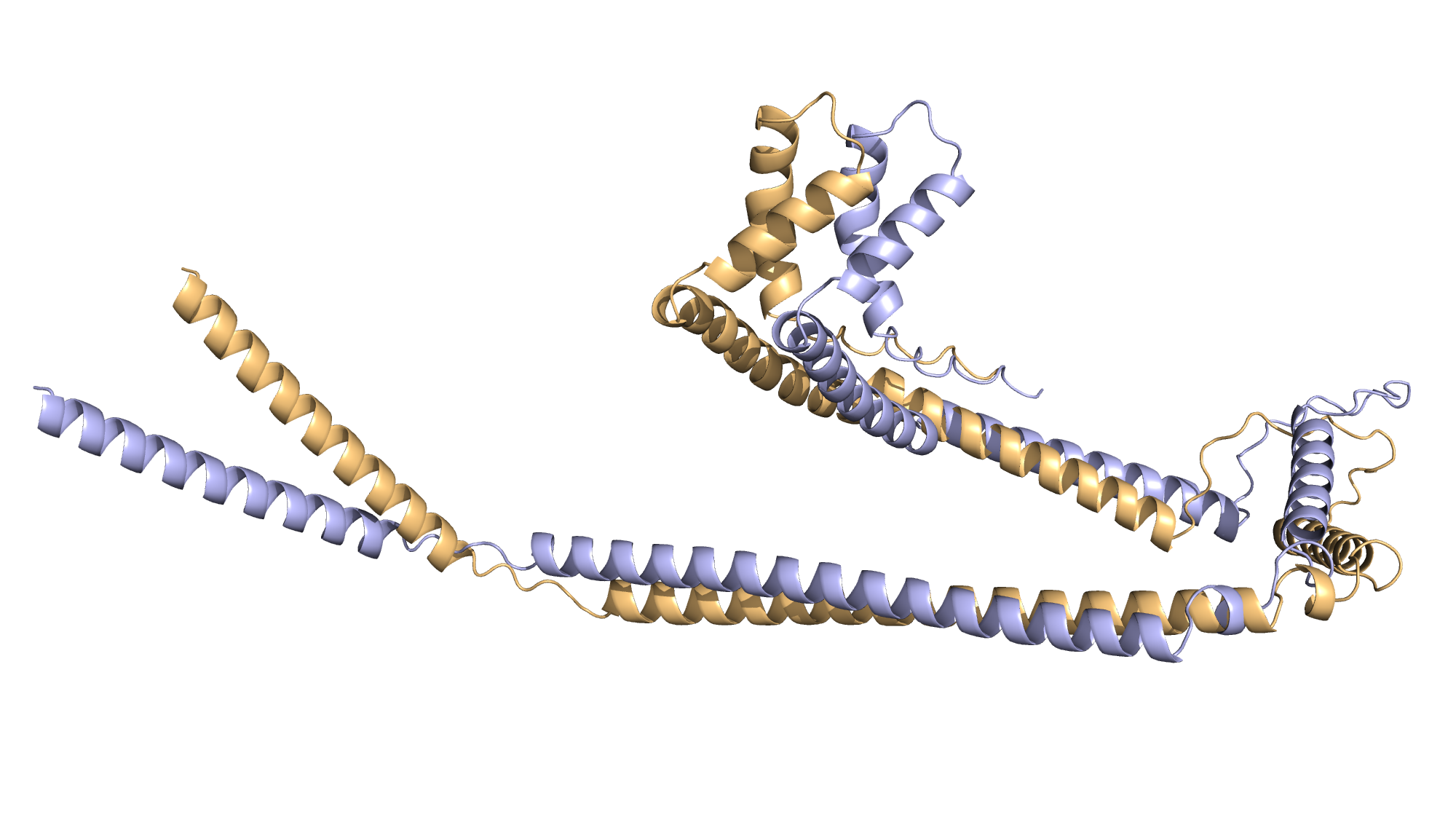

**Figure S1.4:** Visual comparison for SMARCE1_58-315_ (C_α_ RMSD 12.5 Å)

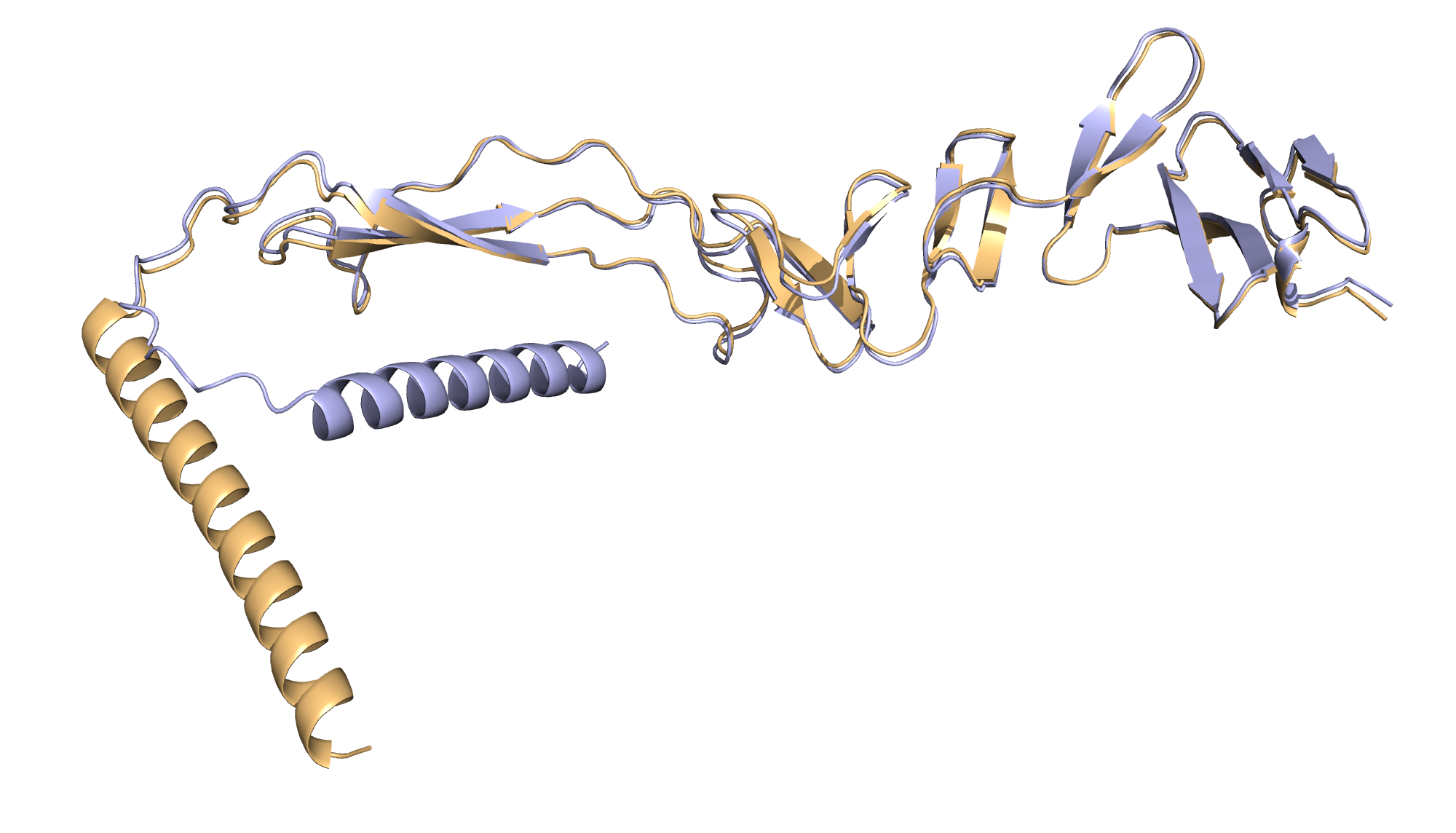

**Figure S1.5:** Visual comparison for RSPO2_35-243_ (C_α_ RMSD 11.0 Å)

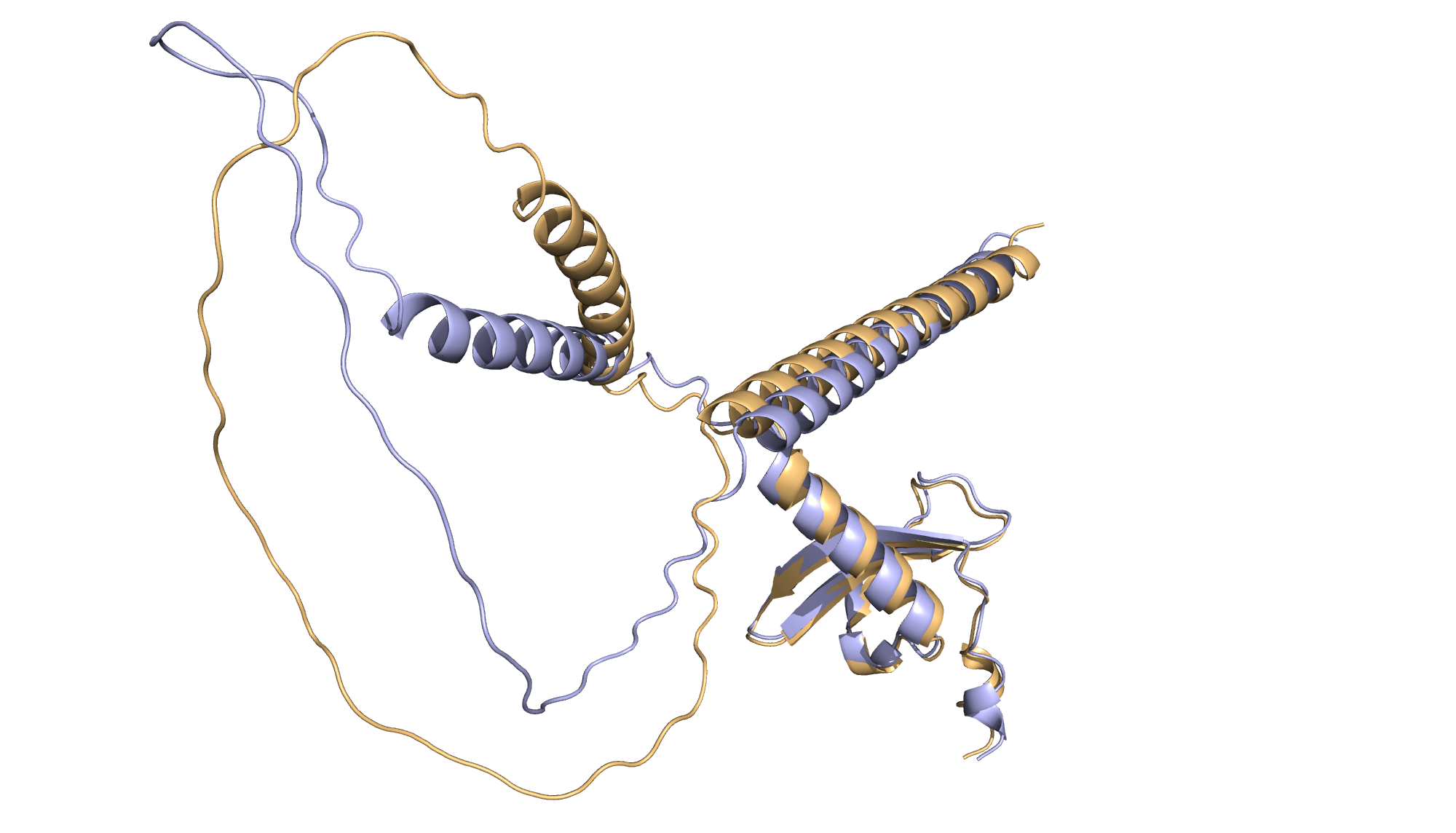

**Figure S1.6:** Visual comparison for CBX8_3-188_ (C_α_ RMSD 9.0 Å)

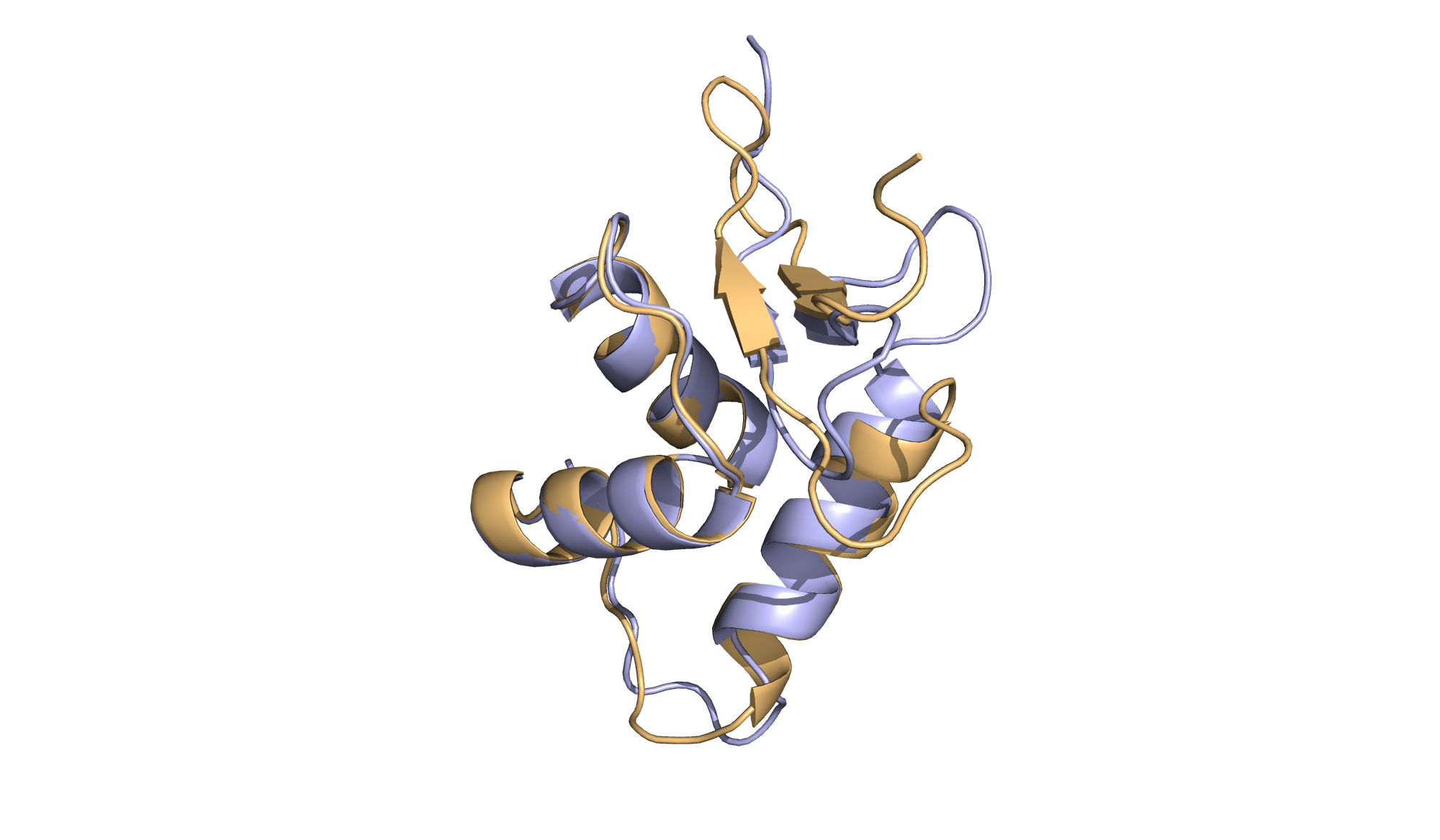

**Figure S1.7:** Visual comparison for ASXL2_8-88_ (C_α_ RMSD 8.6 Å)

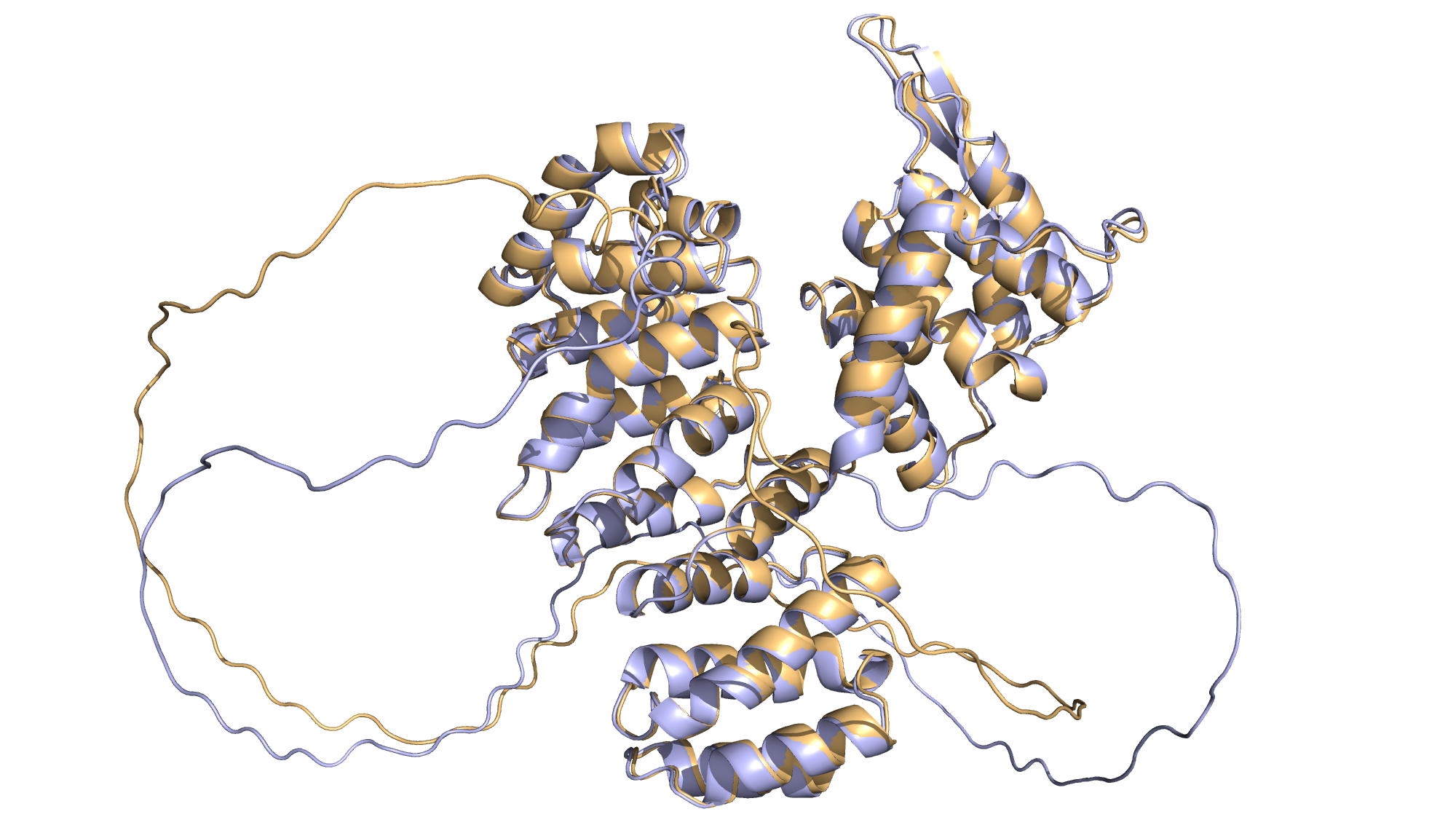

**Figure S1.8:** Visual comparison for FANCE_18-534_ (C_α_ RMSD 7.9 Å)

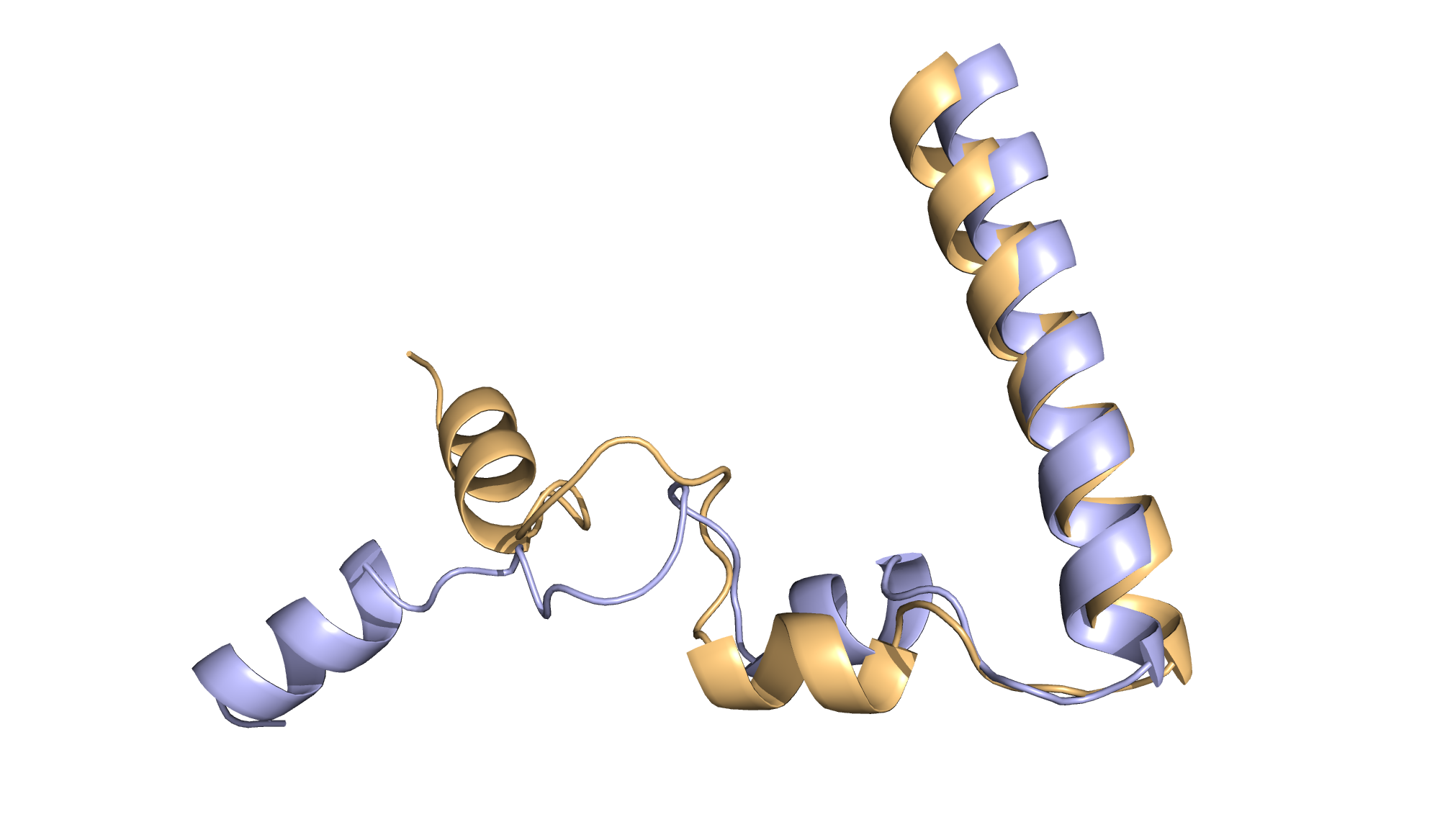

**Figure S1.9:** Visual comparison for NUPR1_24-82_ (C_α_ RMSD 6.8 Å)

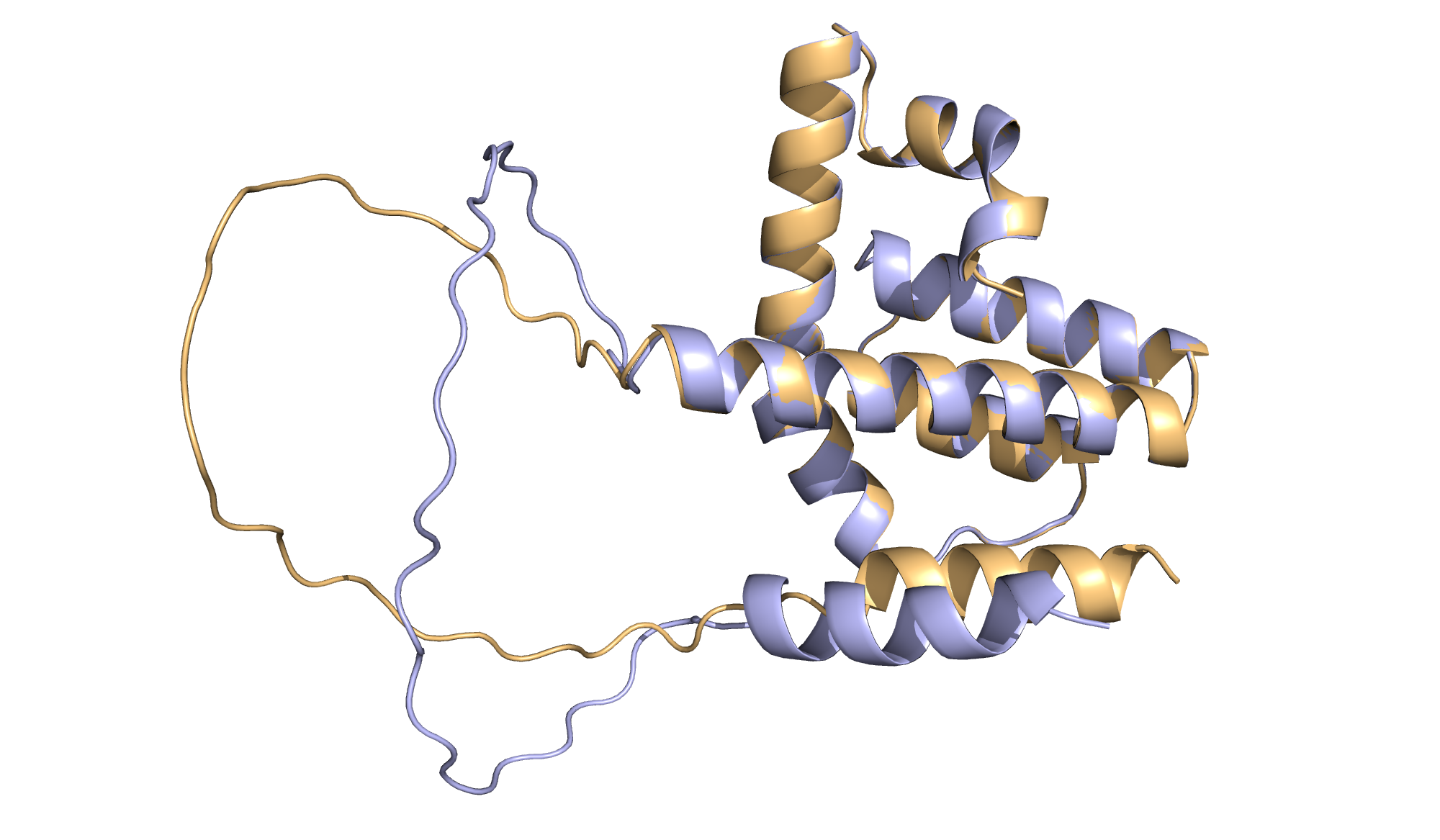

**Figure S1.10:** Visual comparison for CBFA2T3_116-270_ (C_α_ RMSD 6.2 Å)
